# *Pyricularia* Populations are Mostly Host-Specialized with Limited Reciprocal Cross-Infection Between Wheat and Endemic Grasses in Minas Gerais, Brazil

**DOI:** 10.1101/2023.01.20.524950

**Authors:** João P. Ascari, Luis I. Cazón, Mostafa Rahnama, Kurt Lamour, José M. C. Fernandes, Mark L. Farman, Emerson M. Del Ponte

## Abstract

Wheat blast, caused by *Pyricularia oryzae* Triticum (PoT), is an emergent threat to wheat production. Current understanding of the evolution and population biology of the pathogen and epidemiology of the disease has been based on phylogenomic studies that compared the wheat blast pathogen with isolates collected from grasses that were invasive to Brazilian wheat fields. Genetic similarity between isolates from wheat and grasses lead to the conclusion that significant cross-infection occurs, especially on signalgrass (*Urochloa spp*.); and this in turn prompted speculation that its widespread use as forage is a key driver of the disease’s epidemiology. We reanalyzed data from those studies and found that all but one of the isolates from non-wheat hosts were members of PoT and the related *Lolium*-adapted lineage (PoL1), which meant that the *Pyricularia* populations typically found on endemic grasses had not yet been sampled. To address this shortcoming, we performed a comprehensive sampling of blast lesions in wheat crops and endemic grasses found in and away from wheat fields in Minas Gerais. A total 1,368 diseased samples were collected (976 leaves of wheat and grasses and 392 wheat heads) which yielded a working collection of 564 *Pyricularia* isolates. We show that, contrary to earlier implications, PoT was rarely found on endemic grasses and, conversely, members of grass-adapted populations were rarely found on wheat. Instead, most populations were host-specialized with constituent isolates usually grouping according to their host-of-origin. With regard to the dominant role proposed for signalgrass in wheat blast epidemiology, we found only one PoT member in 67 isolates collected from signalgrass grown away from wheat fields, and only three members of *Urochloa*-adapted populations among hundreds of isolates from wheat. Cross-inoculation assays on wheat and a signalgrass used in pastures (*U. brizantha*) suggested that the limited cross-infection observed in the field may be due to innate compatibility differences. Whether or not the observed level of cross-infection would be sufficient to provide an inoculum reservoir, or serve as a bridge between wheat growing regions, is questionable and, therefore, deserves further investigation.

## Introduction

The ascomycete *Pyricularia oryzae*, one of the most studied and economically important fungal plant pathogens worldwide (Dean et al. 2012), is the cause of diseases in commercial crops including rice blast (Valent and Chumley, 1991), wheat blast (Couch and Kohn, 2002) and gray leaf spot in annual and perennial ryegrasses (Farman, 2002). The disease is a current threat to the cultivation of wheat on three continents: South America, Asia, and Africa. First reported in 1985 in the state of Paraná, Brazil (Igarashi et al. 1986), wheat blast spread to all major Brazilian wheat-producing regions (Ceresini et al. 2018; Goulart et al. 1990), and to neighboring countries including Paraguay, Bolivia, and Argentina (Barea and Toledo, 1996; Cabrera and Gutierrez, 2007; Cazal-Martínez et al. 2021). International attention has been raised after the discovery of wheat blast in Bangladesh, south Asia (Malaker et al. 2016) and, more recently, in Zambia, Eastern Africa (Tembo et al. 2020).

Wheat blast epidemics occur more frequently in the tropics where significant yield losses have been associated more often with symptoms on the heads than on the leaves (Cruz and Valent, 2017). In fact, leaf blast sporadically occurs in Brazilian wheat fields when warm and wet weather during the early season might favor infection and inoculum build-up on young leaves (Cruz et al. 2015). The first detailed report of yield losses due to wheat blast was estimated at around 27% in Brazil (Goulart et al. 1990), but greater losses, nearing 100%, have also been reported (Coelho et al. 2016; Dianese et al. 2021; Santos et al. 2022; Goulart and Paiva, 2000; Trindade et al. 2006). In the first major epidemics in Bangladesh, in 2016, the disease caused losses in more than 15,000 ha, which resulted in complete destruction of some affected fields (Islam et al. 2016; Malaker et al. 2016).

*P. oryzae* is a species comprising more than 30 sub-populations that are delimited primarily based on host affinity, although there is evidence that significant gene flow and admixture has occurred amongst them (Gladieux et al. 2018; Valent et al. 2019). These sub-populations, also known as pathotypes (Ou, 1980), or lineages (phylogenetically distinct groups) (Talbot et al. 1993), are normally specialized on one particular host (Gladieux et al. 2018), with some exhibiting fairly strict host-specificity with almost no evidence of cross-infection in nature. These include the lineages found on *Oryza (P. oryzae Oryza*, PoO), *Setaria* (PoS), *Stenotaphrum* (PoSt), and *Eleusine* (PoE) (Gladieux et al. 2018; Latorre et al. 2020). On the other hand, the *Triticum*-specialized lineage (PoT), and the related *Lolium* pathogens (PoL1), among others, should be more accurately defined as host-specialized because, although these are mostly found in association with their eponymous hosts, they can also be found on other Poaceae species (Kato et al. 2000; Tosa et al. 2004; Tosa and Chuma, 2014; Urashima et al. 1993). The ability of the wheat blast pathogen to infect additional cereal crops, as well as forage and turf grasses, is important when thinking about disease development and epidemic spread. This is because alternative hosts often occur in proximity to wheat fields and may occupy large geographical areas, either due to their invasive nature, or their widespread use as forage.

There have been some studies on the population structure and genetic diversity of wheat blast in Brazil and its relationship to isolates found on surrounding grasses (Castroagudín et al. 2017; Castroagudín et al. 2016; Maciel et al. 2014, 2023). In 2012, Ceresini and coworkers collected a large number of wheat blast isolates from more than ten locations in seven states of Brazil. At the same time, they sampled *P. oryzae* from grasses bordering the wheat fields, as well as isolates from rice production areas (Castroagudín et al. 2016). These studies revealed a strong phylogenetic relationship between isolates from wheat and certain grasses, and a distinct separation from PoO, which led to the proposition of a new species, *Pyricularia graminis tritici* (Pygt) (Castroagudín et al. 2016). Subsequent studies implied there was significant evidence of gene flow between the wheat blast and grass-infecting isolates (Castroagudín et al. 2017), prompting speculation that wheat blast undergoes mating on endemic grass species, thereby increasing genetic diversity within the blast population (Ceresini et al. 2018, 2019). Lastly, it was suggested that isolates causing wheat blast showed a particularly close taxonomic affinity with isolates from signalgrass (*Urochloa* spp.) - a widely grown forage crop in Brazil - promoting the hypotheses that wheat blast evolved via a host jump from *Urochloa* (Stukenbrock and McDonald, 2008); and that *Urochloa* serves as a key inoculum reservoir, and a “bridge” facilitating gene flow between separate wheat growing regions (Ceresini et al. 2018, 2019).

However, as noted in the accompanying paper (Farman et al. 2022), when the fungal isolates used by Ceresini and colleagues were analyzed in a broader phylogenetic framework, this revealed that the foregoing studies had not actually sampled the endemic grass-infecting populations because the isolates from grasses were PoT and PoL1 lineage members - probably from opportunistic infections on grasses invasive to wheat crops. Moreover, a preliminary survey based on genome sequencing of a sample of grass-infecting isolates collected at varying distances away from wheat fields suggested that PoT is rarely found on endemic grasses. This latter finding motivated the present study where we sought to characterize the endemic grass-infecting populations in the Cerrado region of Minas Gerais (MG) state with the specific goals of testing the following hypotheses: 1) Infection of endemic grasses, and especially signalgrass, by the wheat-infecting (PoT) lineage is mostly restricted to plants in and around wheat fields, where wheat blast inoculum densities are highest; 2) fungal isolates that typically infect native grasses are rarely found on wheat; and 3) signalgrass/wheat does not support effective colonization of plant tissue by PoT/non-PoT lineage members. To test these hypotheses, we comprehensively sampled *P. oryzae* from wheat fields and from grasses growing at varying distances from wheat-growing locations. PCR assays and genotyping-by-sequencing were then performed to identify isolates down to species and lineage levels, thereby providing an accurate insight into the relationship between fungal populations infecting wheat and grasses. A particular focus was placed on populations infecting signalgrass to re-evaluate the hypothesis that they play a major role in wheat blast epidemiology.

## Materials and Methods

### Study area and sampling

Surveys were conducted in wheat-growing regions and natural landscapes of MG state during the 2018 and 2019 growing seasons. While wheat blast was found only on the heads of wheat crops in 2018 (n = 4 wheat fields), it occurred both on leaves and heads of wheat in 2019 (n = 11 wheat fields). The sampling target and design varied according to the timing of sampling and whether the site was a wheat or non-wheat area (Fig. 1). The pre-season sampling in mid-February (summer season in MG) targeted grass weed hosts which were collected randomly by visiting natural landscapes along roadsides and in off-season wheat areas (Fig. S1). Each sample comprised five to ten leaves, which were placed into paper bags and placed at room temperature (23°C ±4°C) to dry for one week before being stored at 10°C. Mid-season sampling in mid to late May (Fall season) focused on collecting: a) blast-symptomatic leaves (only in 2019) and heads (2018 and 2019) in wheat fields and b) blast-symptomatic leaves of grass weeds either within or near to wheat growing areas. For the wheat blast samples, five to 10 (depending on field size) 50-m transects placed 200 m apart were randomly defined. At least one sample (five to ten leaves or five heads) was collected at each transect, similar to a previous study (Maciel et al. 2014). Weed species were identified morphologically based on the literature (taxonomic guides) (Lorenzi, 2014). The wheat varieties could not be identified. All natural landscapes, wheat commercial fields, and individual plants in the field were photographed using a smartphone camera (72 dpi resolution). In the laboratory, photographs of the symptoms, on leaves or heads (in the case of wheat), were obtained using a smartphone camera and a digital magnifying miniscope (10X, 96 dpi resolution) (Fig. 2S).

**Fig. 1.**
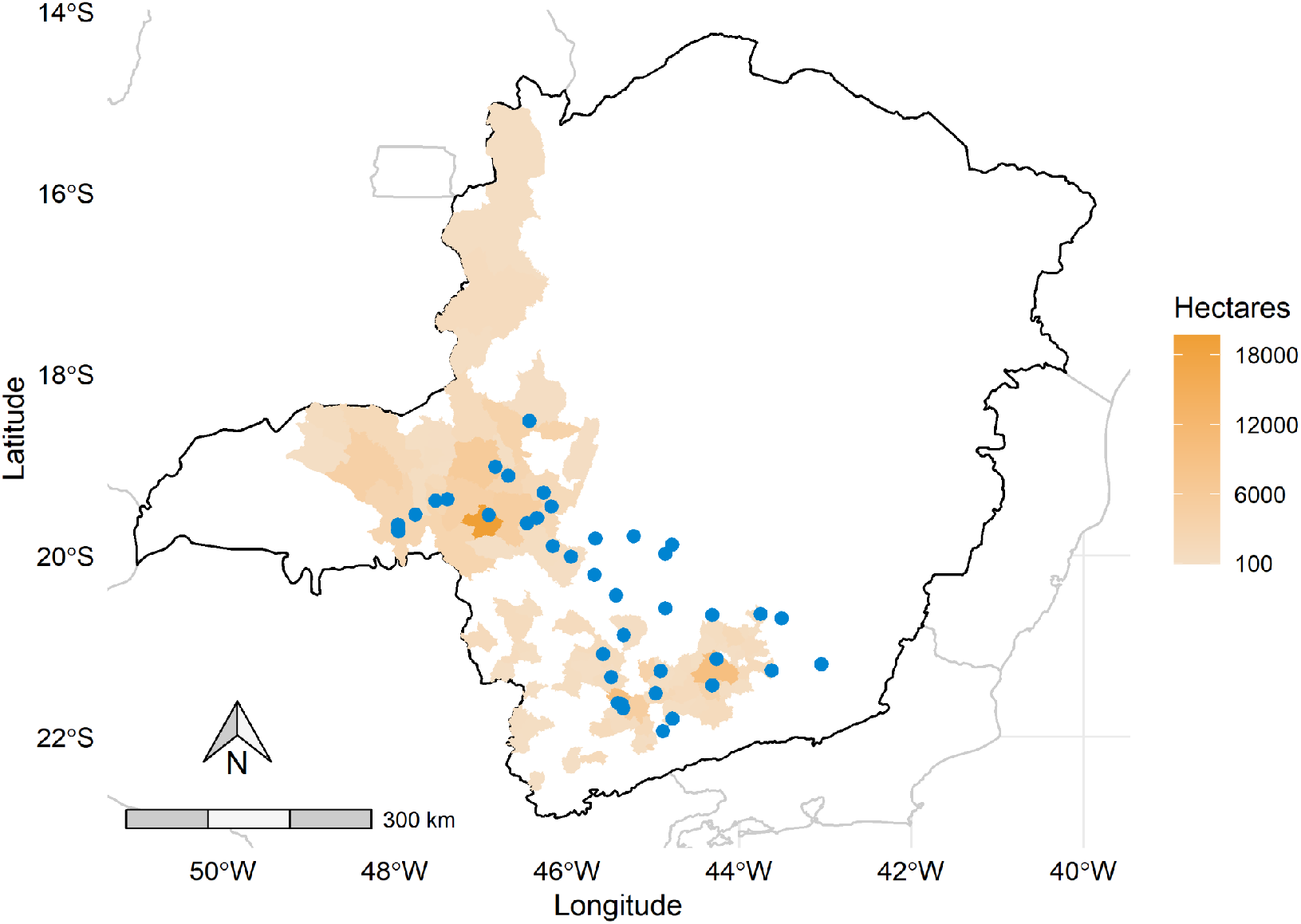
Map of Minas Gerais (MG) state, Brazil, depicting wheat area (in hectares) planted per municipality (color gradient) and the locations of the sampling sites where blast-symptomatic Poaceae (wheat and grasses) were collected (dots). Source: (IBGE, 2017).

**Fig. 2.**
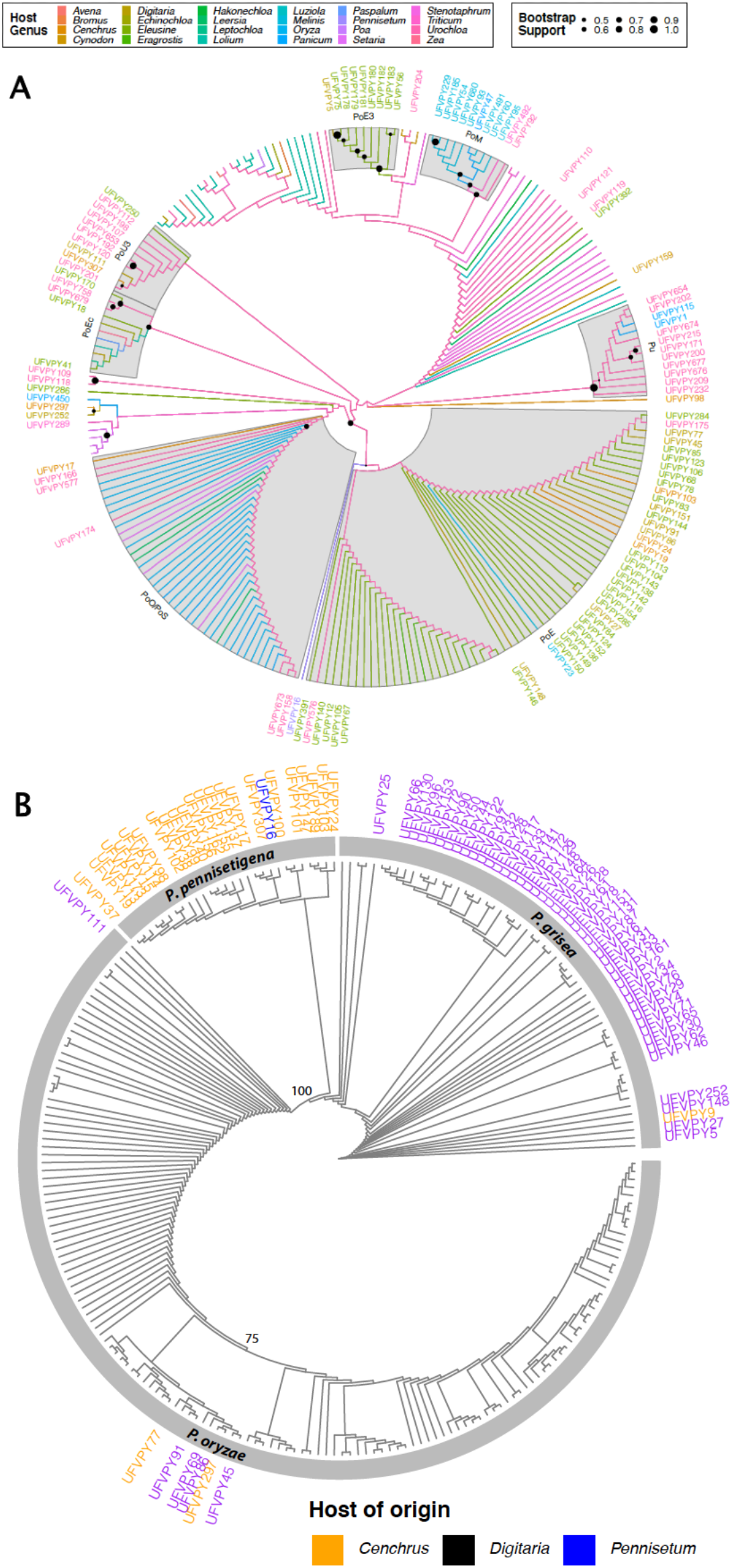
Maximum likelihood trees used for lineage/species assignment of *Pyricularia* isolates collected from grasses in MG. **A**, CH7BAC9 tree showing phylogenetic placement of *P. oryzae*/*P. urashimae* isolates. Tip labels are provided only for isolates collected in this study and are colored according to the host-of-origin, as are the branches for the unlabeled reference isolate. Phylogenetic lineages showing obvious host-specialization are highlighted with gray boxes and named according to primary host as follows: PoEc, *Echinochloa*; PoE/PoE3, *Eleusine*, PoM, *Melinis*; PoO/PoS *Oryza/Setari*a; PoU3, *Urochloa*; and Pu, *Pyricularia urashimae*. Nodes with bootstrap support ≥ 0.5 are highlighted with proportionally-sized circles. The tree was drawn using ggtree with the branch.length = “none” option, with the intention to show grouping patterns among isolates from the same host. No inferences can be drawn from branch lengths. **B**, MPG1 tree showing phylogenetic placement of *Pyricularia* isolates collected from *Cenchrus, Digitaria, Panicum* and *Urochloa*. For clarity, tip labels are shown only for the isolates from MG, and their prefixes are in lowercase and abbreviated. Bootstrap values are provided on key nodes. Tip labels are colored according to host-of-origin. The species designation for each phylogenetic clade is also shown.

### Culturing, purification and storage

Wheat heads (one per sample) and leaves (five per sample) of wheat or grass weeds, were cut into small pieces and placed within a 9 cm-plastic dish filled with moistened filter paper, and incubated for 24 h at 25°C ±5 under a 12/12 h photoperiod (light/darkness) to induce sporulation (Urashima et al. 2017). Under the stereomicroscope light, conidiophores and associated sparkling crystal-clear spore mass on leaf and head-rachis could be visualized. A sterilized sealed Pasteur pipette was scraped over the sporulating mass and streaked across the water agar supplemented with chloramphenicol and streptomycin, each at 100 μg/ml. Plates were incubated at 25±5°C for 24 h (12/12 h fluorescent light/darkness) (Farman et al. 2017; Gupta et al. 2020). For each culture, a single biseptate, pyriform conidium (Klaubauf et al. 2014; Murata et al. 2014) with a visible germ tube was transferred to oatmeal agar (OA) (30 g oats, 20 g agar, 1 L distilled water), and pieces of sterilized filter paper (10 mm x 0.4 mm) were placed nearby. The dishes were incubated as above for 7 d until the mycelium fully covered the filter paper. The papers were then transferred to a new Petri plate filled with blue silica crystals and left to dry at room temperature (25°C±5) for 5 d. Dried paper pieces were transferred to a 2 ml-microtube half-filled with fresh sterile blue silica and stored in a -10°C freezer (Farman et al. 2017; Farman, 2002; Gupta et al. 2020). Isolates were stored in duplicates as a backup of the entire collection.

### Growth of *Pyricularia* spp. and DNA extraction

A single filter paper of each isolate was placed on a potato dextrose agar and incubated at 25±5°C under 12/12 h photoperiod (fluorescent light/darkness). A 6 mm mycelial block from a 5-day-old colony was then transferred to a 50 mL falcon tube filled with 20 mL of liquid complete medium (6 g casamino-acids, 6 g yeast extract, and 10 g sucrose per 1 liter). The tubes were shaken for 7 days at 150 rpm under room temperature (23-26°C) and ambient light. The mycelium was recovered through two layers of cheesecloth and let to dry at an ambient temperature for 3 h, and freeze-dried in 2 mL microtubes for 24 h (M. Farman et al. 2017; Urashima et al. 2017) using a CoolSafe Freeze Dryer (SCANVAC). The mycelium ball was manually crushed against the microtube wall until it formed a powder, which was then resuspended in 1 mL lysis buffer (100 mM Tris-HCl, pH8; 0.5 M NaCl, 10 mM EDTA; 1 % SDS) and heated 65°C for 30 min. Adding 700 μl phenol:chloroform:isoamyl alcohol (25:24:1) and heated 65°C for 30 min. Subsequently, centrifuged at 14000 rpm for 15 min, and carefully transferred 0.8 μl of aqueous phase to a new identified microtube, where was added 450 μl of cool isopropanol and centrifuged at 14,000 rpm for 10 min to pellet the DNA. The supernatant was carefully discarded and the pellet was washed with 1 mL of 70% ethanol, and re-pelleted by centrifuging for 5 min at 14,000 rpm. The supernatant was discarded and the DNA was dried at room temperature for 60 min, redissolved in 100 μl TE + 2 μl RNAse A (1 μg/ml), and stored at 4°C overnight, before being placed in the -20°C freezer (Farman et al. 2017). The DNA concentration was estimated using a spectrophotometer NanoDrop 2000 (Thermo Scientific™) and adjusted to 100 ng/μl using TE buffer.

### PCR assays targeting *P. oryzae*

The entire collection of 572 isolates was first screened by using PCR to amplify the CH7-BAC9 locus, which is present in *P. oryzae* (Po) but absent in other *Pyricularia* (non-Po), including *P. grisea, P. pennisetigena*, or *P. urashimae* (Couch et al. 2005). The assays were performed using 1 μl of genomic DNA (100 ng/μl) and primer concentrations of 10 μM, with the GoTaq® Colorless Master Mix, according to the manufacturer’s specifications (Promega). Reactions were carried out in a MyGene™ thermal cycler (Model MG96G), with the following parameters: an initial denaturation at 95° for 8 min, followed by 35 cycles of 95°C for 15 sec, 55°C for 20 sec, 72°C for 60 sec, and a final extension at 72°C for 5 min (Couch et al. 2005). To confirm the accuracy of CH7-BAC9 for discriminating Po from non-Po, the MPG1 locus was amplified and sequenced for all negatives and select positive ones. The sequence of this gene was used in phylogeny analysis to identify *Pyricularia* at the species level (Couch et al. 2005). PCR assays were performed with 1 μl of genomic DNA (100 ng/μl) using the same GoTaq® Mix and thermal cycler. Amplification conditions for MPG1 were as follows: initial denaturation at 95° for 8 min, followed by 35 cycles of 95°C for 15 sec, 55°C for 20 sec, 72°C for 60 sec, and a final extension at 72°C for 5 min (Couch et al. 2005). PCR products were sequenced.

### PCR assays targeting *P. oryzae* Triticum pathotype

To distinguish *P. oryzae* Triticum lineage members (PoT) from non-PoT, we used the MoT3 primer set (F: GTCGTCATCAACGTGACCAG; R: ACTTGACCCAAGCCTCGAAT) that yields a 362 bp amplicon (Pieck et al. 2017). For C17 diagnostics (Thierry et al. 2020), we designed a new primer set for a modified (standard PCR) assay that identifies PoT based on the positive amplification of a 500 bp fragment (F: GAGGAAGATCAAGTAAGTGG; R: GGTAGATGTCATGATTTCAC). Here, it is important to note that while these two loci were selected for the specific purpose of identifying PoT (MoT), neither is truly diagnostic because both loci were contributed to the PoT lineage via admixture (Rahnama et al. 2021). MoT3 was donated by a *Urochloa* pathogen from the PoU3 (subgroup of PoSt) lineage and, therefore, tests positive with certain isolates from *Urochloa*. Likewise, C17 was contributed to PoT by a lineage that is related to rice pathogens, but has not yet been sampled from the field (“PoX”). For this reason, tentative lineage designations were made according to a specific schema (Table 1) and, where necessary, sequencing of the CH7-BAC9 and MPG1 loci, and genotyping by sequencing were performed to validate the assignments.

**TABLE 1.** Identification of *P. oryzae/P. urashimae* lineages based on MoT3/C17 test results

| lineage | MoT3 | C17 |
| --- | --- | --- |
| PoT | + | + |
| PoU3, Pu<sup>a</sup>, (PoT)<sup>b</sup> | + | - |
| PoT, PoX<sup>c</sup> | - | + |
| PoL, PoM, others<sup>d</sup> | - | - |

<sup>a</sup> *Pyricularia urashimae*
<sup>b</sup> Eventual detection of PoT (*Pyricularia oryzae* *Triticum* lineage) members with this pattern is predicted
<sup>c</sup> Definitive identification of the lineage requires sequencing of additional loci
<sup>d</sup> Definitive identification of the lineage requires sequencing of additional loci or genome sequencing

Assays were performed using 10 ng template and the same GoTaq® mix. Cycling conditions were as follows: MoT - initial denaturation at 95°C for 8 min, followed by 35 cycles of 95°C for 15 sec, 55°C for 20 sec, 72°C for 60 sec, and a final extension at 72°C for 5 min; C17 - initial denaturation at 94°C for 2 min, 35 cycles of 95°C for 10 sec, 54°C for 30 sec, 72°C for 30 sec, and a final extension at 72°C for 5 min. PCR products were fractionated by electrophoresis in a 1%-agarose gel for 100 min at 80 Volts, 100 mA, and 80 watts, using a 1 Kb DNA ladder (Cellco®). The DNA was stained with GelRed® and the gel was visualized and photographed under ultraviolet (UV) light.

### Lineage assignment

Consensus lineage assignments were made using different criteria and took into account the host-of-origin, established patterns of sequence distribution among the different host-specialized lineages, as well as known population structure. For example, if an isolate came from a host that typically harbors one or more specific lineages, and those lineages have not yet shown any evidence of admixture, sequence data for a single marker allowed a confident assignment. For others - especially isolates from lineages with higher than normal cross-infection behavior, or known admixture - multi-locus genotyping was necessary.

### Cross-inoculation assays

A subcollection of 20 strains isolated from wheat (n = 11, being all PoT) or signalgrass (n = 9, being six non-PoT [three PoU and two Pu] and three PoT) was studied with regards to aggressiveness towards leaves and heads of two wheat cultivars (BR 18-Terena and BRS Guamirim) and leaves of one *Urochloa brizantha* cultivar (cv. Marandu) (Table 2). Among the PoT isolates, a reference isolate (16MoT001), used as standard for aggressiveness in screening for host resistance (Cruppe et al. 2020), was included for comparison. The inoculations on the leaves were conducted on 35-day-old plants exhibiting three to four completely expanded leaves, growth stage 15 (Zadoks et al. 1974). Inoculations on the heads were performed in 60-day-old plants at early anthesis, growth stage 60 (Zadoks et al. 1974). Each experiment (inoculation on leaves or heads) was conducted twice under greenhouse conditions between March and September 2020.

**TABLE 2.** Information for isolates obtained from wheat blast (PoT = 14 isolates) or signalgrass blast (PoT = 3; non-POT = 6; species/lineage as designated) used in replicated cross-inoculation experiments.

| Host of Origin | Municipality | Collection date | Code<sup>b</sup> | ID<sup>c</sup> |
| --- | --- | --- | --- | --- |
| *Urochloa brizantha* | Patos de Minas | Feb. 2018 | 108 | PoU3 |
| *U. brizantha* | Patos de Minas | Feb. 2018 | 110 | PoU3 |
| *U. brizantha* | Patos de Minas | Feb. 2018 | 112 | PoU3 |
| *U. brizantha* | Uberaba | May 2018 | 166 | PoU4 |
| *U. brizantha* | Formiga | Feb. 2019 | 209 | Pu |
| *U. brizantha* | Catas Altas da Noruega | Feb. 2019 | 656 | Pu |
| *U. brizantha* | Madre de Deus | May 2019 | 742 | PoT |
| *U. brizantha* | Madre de Deus | May 2019 | 758 | PoT |
| *U. humidicola* | São Gonçalo do Pará | Feb. 2019 | 213 | PoT |
| *Triticum aestivum* | Patos de Minas | May 2018 | 167 | PoT |
| *T. aestivum* | Ibiá | May 2019 | 238 | PoT |
| *T. aestivum* | Ibiá | May 2019 | 239 | PoT |
| *T. aestivum* | Uberaba | May 2019 | 367 | PoT |
| *T. aestivum* | Santa Juliana | May 2019 | 375 | PoT |
| *T. aestivum* | Patrocínio | May 2019 | 376 | PoT |
| *T. aestivum* | Boa Esperança | May 2019 | 309 | PoT |
| *T. aestivum* | Boa Esperança | May 2019 | 311 | PoT |
| *T. aestivum* | Madre de Deus | May 2019 | 604 | PoT |
| *T. aestivum* | Boa Esperança | May 2019 | 813 | PoT |
| *T. aestivum* | Passo Fundo, RS | 2019 | MoT01 | PoT |

<sup>a</sup> Sampling distance from wheat areas. Within (in-field); Nearby (< 1 km) and Away (> 50km).
<sup>b</sup> Prefix UFVPY
<sup>c</sup> The separation between PoT and non-PoT isolates was performed using MoT3 and C17 primers in a PCR assay. The non-PoT lineages were identified based on genotyping by sequencing data. PoU = *Pyricularia oryzae* lineage *Urochloa*; and Pu = *Pyricularia urashimae*

#### Inoculum production

For each isolate, a piece of filter paper containing the fungus was removed from the -10°C storage and re-activated on Potato Dextrose Agar (PDA). A 5-day-old mycelial plug was transferred to oatmeal-agar (OA) (replicated in five 9 cm-dishes per isolate). The fungus was cultured for seven days. To induce fungal sporulation, plates were scraped out using a Drigalski spatula and 5 ml of sterilized-distilled water. The dishes were incubated for a further seven days. Spores were harvested by adding 10 ml of distilled-sterilized water amended with 0.01% Tween-20, and carefully scraped using a Drigalski spatula. Spore suspension was filtered through two layers of cheesecloth. Spore concentration was adjusted to 1×10^5^ spores/mL using a Neubauer counting chamber. PDA and OA dishes were both supplemented with chloramphenicol and streptomycin at 100 μg/ml. Incubation was performed in a grown chamber with controlled temperature of 25°C (±2°), and photoperiod of 12/12 hours (fluorescent light/darkness) (Cruz et al. 2016; Urashima et al. 2017).

#### Plant growth conditions

The plants were sown in 2-L plastic pots filled with substrate (Tropstrato - Vida Verde) which was a mixture of pine bark, peat, and expanded vermiculite. Basal fertilization was performed with monoammonium phosphate (12% N and 50% P_2_O_5_). The number of plants per pot was reduced to eight and ten for wheat and signalgrass, respectively. Plants were kept in the greenhouse under controlled environmental conditions (±11 hour of light and 25°C ±4°C) and watered daily until inoculation time.

Side-dressing fertilization were conducted weekly adding to each pot 30ml of nutritive solution prepared with 6.4mg/L KCl, 3.48mg/L K_2_SO_4_, 5.01mg/L MgSO_4_.7H_2_O, 2.03mg/L (NH_2_)2CO, 0.009mg/L NH_4_MO_7_O_24_.4H_2_O, 0.054mg/L H_3_BO_3_, 0.222mg/L ZnSO_4_.7H_2_O, 0.058mg/L CuSO_4_.5H_2_O, 0.137mg/L MnCl_2_.4H_2_O, 0.27g/L FeSO_4_.7H_2_O and 0.37g/L disodium-EDTA prepared with distilled water (Xavier Filha et al. 2011).

#### Inoculation procedures

Plants (leaves or heads) on each pot were sprayed-inoculated (15 mL) with the spore suspension using a 0.5L manual plastic sprayer (Guarany® - Gifor). The plants were placed in the dark within a chamber adjusted to 25°C (±2%) and humidity >90% during 20 hours. The potted plants were moved to a growth chamber with controlled temperature at 28°C (±2°), humidity >80%, and 12/12 hours of fluorescent light/darkness during seven days, until performing the disease assessments.

#### Disease assessment and data analysis

The assessment of leaf blast severity (percentage area affected) in wheat and signalgrass, and severity on wheat heads (percent of spikelets with symptoms), was conducted seven days post-inoculation (dpi). Severity on the leaves was measured on ten leaves randomly selected from each pot. These were removed from the plant and imaged against a white background, using a flatbed scanner (HP - LaserJet M1132 MFP) at 600-dpi resolution and JPEG file format. Images were analyzed in ImageJ (Schneider et al. 2012) to threshold the symptomatic and asymptomatic area, and then calculate severity (% symptomatic area). The severity on wheat heads was assessed visually. The means and 95% confidence interval of the percent severity of leaf and head blast were estimated for each isolate after pooling data from two replicates of the experiment.

## Data and code availability

Data and custom R codes (R Core Team 2022) used for data analyses are available at https://github.com/emdelponte/paper-wheat-blast-MG

## Results

### Recovery of *Pyricularia* from blast-like lesions

A large collection of isolates from wheat and grasses, both nearby and away (dozens to hundreds of km) from wheat crops, were obtained during the two-year, multi-location, survey conducted across the state of MG. A total of four surveys were conducted - prior to and during the wheat growing seasons - in 2018 and 2019, which experienced typical and severe wheat blast outbreaks, respectively. Poaceae with blast-like symptoms (diamond-shaped lesions) were sampled during the visits, but with a focus on signalgrass (*Urochloa* spp.) growing near to wheat fields, and in natural landscapes farther away. A total of 1,368 diseased samples were collected (976 for leaves and 392 for wheat heads) from 20 Poaceae genera (31 species) (Table 3). Symptomatic plants collected at the non-wheat regions comprised mostly weeds and included *Cenchrus, Cynodon, Digitaria, Eleusine, Hordeum, Melinis*, and *Panicum*. Symptoms were found on six species of signalgrass (*U. brizantha, U. humidicola, U. plantaginea, U. ruziziensis, U. arrecta, U. decumbens*), with *U. brizantha* being the most prevalent. In total, pure cultures that were morphologically similar to *Pyricularia* spp. were successfully recovered from approximately 68% of samples, resulting in a total of 932 monoconidial isolates (Table 3).

**TABLE 3.** Summary information for the total number of blast-infected plant samples for each of 31 Poaceae species, including wheat, from where *Pyricularia* sp. isolates were obtained during four visits to both wheat-producing regions (Triângulo Mineiro and Centro-Sul de Minas) and natural landscapes during summer (February, wheat off-season) and fall (May, wheat-growing season) 2018 and 2019, MG, Brazil.

| Poaceae species | N. of plant samples |  | N. of *Pyricularia* spp. isolates <sup>c</sup> |
| --- | --- | --- | --- |
|  | Total <sup>a</sup> | Blast-infected <sup>b</sup> |  |
| *Andropogon virginicus* | 4 | - | - |
| *Cenchrus echinatus* | 21 | 15 | 25 |
| *Chloris polydactyla* | 4 | - | - |
| *Chrysopogon zizanioides* | 4 | - | - |
| *Cynodon dactylon* | 6 | 1 | 1 |
| *C. dictyoneura* | 1 | - | - |
| *C. plectostachyus* | 1 | 1 | 1 |
| *Cyperus rotundus* | 12 | - | - |
| *Digitaria horizontalis* | 37 | 14 | 21 |
| *D. insularis* | 46 | 15 | 20 |
| *D. sanguinalis* | 61 | 21 | 31 |
| *Echinochloa colonum* | 2 | 1 | 1 |
| *Eleusine indica* | 50 | 27 | 42 |
| *Eragrostis ciliaris* | 6 | - | - |
| *E. pilosa* | 1 | - | - |
| *Imperata brasiliensis* | 2 | - | - |
| *Melinis minutiflora* | 31 | - | - |
| *Melinis roseum* | 20 | 16 | 28 |
| *Panicum maximum* | 127 | 10 | 19 |
| *P. miliaceum* | 3 | - | - |
| *Paspalum notatum* | 4 | - | - |
| *Pennisetum sp.* | 28 | 2 | 2 |
| *Setaria viridis* | 1 | - | - |
| *Sorghum arundinaceum* | 11 | - | - |
| *Triticum aestivum* | 505 | 377 | 670 |
| *Urochloa arrecta* | 1 | - | - |
| *U. brizantha* | 329 | 35 | 59 |
| *U. decumbens* | 2 | - | - |
| *U. humidicola* | 16 | 3 | 7 |
| *U. plantaginea* | 24 | 3 | 3 |
| *U. ruziziensis* | 8 | 1 | 2 |
| Total | 1,368 | 542 | 932 |

<sup>a</sup> Sample composed of 5 to 10 leaves of wheat or Poaceae weeds, or 1 to 3 wheat heads. <sup>b</sup> At least one *Pyricularia* sp. isolate. <sup>c</sup> Number of monoconidial isolates per sample. In cases more than one isolate was obtained from a sample, but not from the same tissue (leaf or head).

### PCR-based diagnostics identified minimal cross-infection between isolates adapted to wheat versus endemic grasses

A subcollection of 564 isolates (including all of those from endemic grasses) were pre-screened using CH7BAC9 PCR (Supplementary Table S1) which yields positive amplification for *P. oryzae* and *P. urashimae* but no products for *P. grisea* or *P. pennisetigena*. Among these, 483 (85.6%) were CH7BAC9-positive and came from 16 plant species, with the predominant hosts being wheat, followed by *Urochloa, Eleusine, Melinis, and Panicum* (Table 4). The CH7BAC9-negative isolates, suspected to be other *Pyricularia* species, mostly came from *Cenchrus echinatus, Digitaria* spp., and *Panicum maximum*.

**TABLE 4.** Summary results of PCR assays in a subcollection of 572 strains from 16 Poaceae hosts recovered of wheat (leaf and head) and grass weeds (leaves) from wheat-producing regions (Triângulo Mineiro and Centro-Sul de Minas) and natural landscapes during summer (February, pre-season) and fall (May, wheat-growing season) 2018 and 2019, MG, Brazil.

| Host of isolation | *Pyricularia* sp. | *P. oryzae* | Lineages |  |
| --- | --- | --- | --- | --- |
|  |  |  | PoT (proximity)<sup>a</sup> | Non-PoT |
| *Cenchrus echinatus* | 20 | 2 | 1 (away) | 1 |
| *Cynodon dactylon* | 1 | 0 | - | - |
| *Cynodon plectostachyus* | 1 | 1 | - | 1 |
| *Digitaria horizontalis* | 21 | 2 | - | 2 |
| *Digitaria insularis* | 20 | 1 | - | 1 |
| *Digitaria sanguinalis* | 18 | 5 | - | 5 |
| *Echinochloa colonum* | 1 | 0 | - | - |
| *Eleusine indica* | 34 | 32 | 2 (nearby) | 30 |
| *Panicum maximum* | 16 | 16 | 1 (nearby) | 15 |
| *Pennisetum sp.* | 2 | 1 | 1 (away) | - |
| *Melinis roseum* | 28 | 28 | 1 (away) | 27 |
| *Triticum aestivum* | 333 | 331 | 324 | 7 |
| Leaf | 127 | 127 | 126 | 1 |
| Head | 206 | 204 | 198 | 6 |
| *Urochloa brizantha* | 57 | 56 | 2 (nearby) | 54 |
| *Urochloa humidicola* | 7 | 7 | 1 (away) | 6 |
| *Urochloa plantaginea* | 3 | 2 | - | 2 |
| *Urochloa ruziziensis* | 2 | 2 | - | 2 |
| Total | 564 | 483 | 329 | 154 |

<sup>a</sup> distance of the isolates obtained from a grass plant relative to wheat fields. Nearby = less than 1km and away = more than 50 km.

Among the 483 *P. oryzae* isolates analyzed by PCR, 313 (64.8%) were identified as PoT based on the successful amplification of both C17 and MoT3. Only nine of these (2.7%) came from non-wheat hosts, with only three coming from *Urochloa* (Table 4). The other grasses found to be harboring PoT were *Cenchrus echinatus* (n = 1 isolate), *Eleusine indica* (n = 2), *Melinis repens* (n = 1), *Panicum maximum* (n = 1), and *Pennisetum sp*. (n = 1). Five of the nine PoT cross-infections on other grasses were for plants collected within, or adjacent to, wheat fields. The four other cases were in locations more than 30 km away from wheat fields, and only one of these remote cross infections was on *Urochloa* (Tables 6).

Sixteen of the isolates from wheat failed to yield amplicons for MoT3 and/or C17 (Table S1). It is well known that certain wheat-infecting PoL1 haplotypes lack both C17 and MoT3 (M. Farman et al. 2017; Pieck et al. 2017; Thierry et al. 2020) and, therefore, eight C17^-^ /MoT3^-^ isolates that came from wheat were tentatively assigned as PoL1, pending further confirmation (see below).

PoT isolates that contain C17 but lack MoT3 are also occasionally found (e.g., Islam *et al*. 2016). However, because C17 and MoT3 actually target loci in native grass-infecting populations (PoX and PoU3, respectively), absences of either marker (or both) yield inconclusive assignments (Table 1). For this reason, we sought to verify equivocal lineage designations by sequencing CH7BAC9 PCR products and/or genotyping-by-sequencing.

CH7BAC9 sequences were obtained for a total of 102 isolates which variously came from *Urochloa* sp. (n = 35), *Eleusine* (n = 42), *Melinis* (n = 8), *Cenchrus* (n = 2), *Panicum* (n = 10) and *Digitaria* (n = 5). These sequences were combined with the broader *P. oryzae* dataset and the phylogenetic relationships between the MG isolates and previously-established lineages were determined using maximum likelihood. This revealed a clear pattern of host specialization because most isolates grouped with lineages whose constituent members usually came from the same host (Fig. 2A, and Table 5). Four different CH7BAC9 alleles were found among the *Urochloa*-infecting isolates with two predominating. One of these matched the PoU3 lineage, while the other was identical to an allele found in a different species - *P. urashimae*. The minor alleles in the *Urochloa* pathogens matched PoM, and PoO/Le/P/S (which are indistinguishable because they all share the same sequence).

**TABLE 5.**
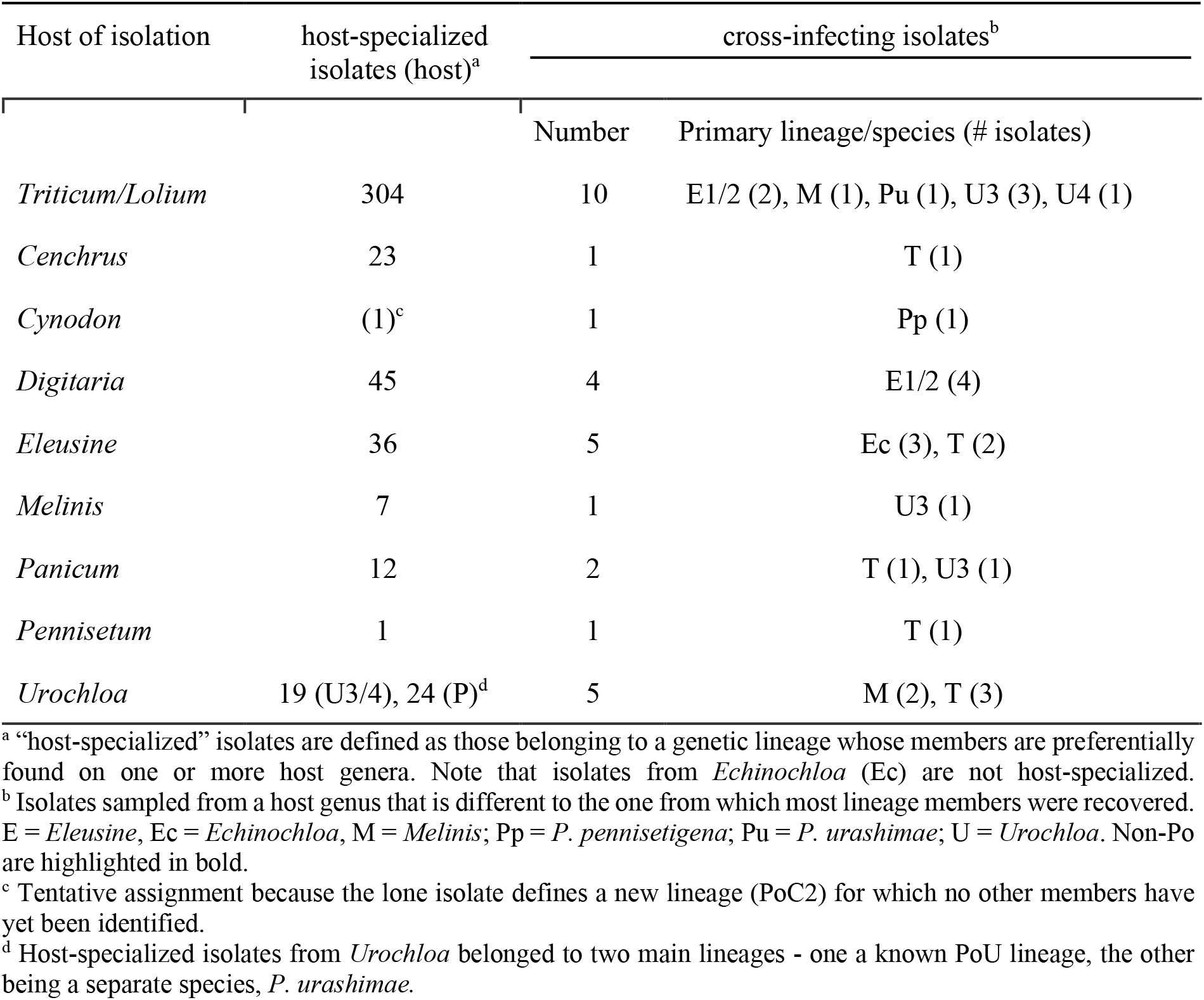
Phylogenetic lineage affiliations of *Pyricularia* isolates obtained from wheat, and endemic/cultivated grasses in MG, Brazil

Most of the isolates from *Eleusine* (80%) grouped with the *Eleusine*-infecting lineages, PoE1/2 (which share the same allele) or PoE3 (28 PoE1/2 and 8 PoE3) (Fig. 2A and Table 5). The only examples of cross-infection on *Eleusine*, were two isolates from PoT, and three from the *P. oryzae Echinochloa* lineage (PoEc), which is represented by isolates from various hosts including *Digitaria, Echinochloa, Lolium, Zea*, and now *Eleusine* (Table 5, Table S1). The CH9BAC9 sequences for the isolates from *Melinis* all grouped with the sequence present in the reference genome of a previous strain from this host, MrJA49 and, therefore, appear to identify a new, phylogenetically-distinct, *Melinis*-adapted lineage.

### Host-specialization in other *Pyricularia* species

No CH7BAC9 PCR products were obtained for most of the isolates (73/84) from *Cenchrus, Digitaria*, and *Pennisetum*, which was consistent with the absence of this locus in the genome assemblies of representative isolates. For the few isolates that did yield amplicons, sequencing revealed that some of these loci were variously related to those found in PoE1/2 (4 isolates from *Digitaria*; 1 from *Cenchrus*, 1 from *Pennisetum*) and PoO (one isolate from *Digitaria*) (Fig. 2A). Given the rarity of cross-species admixture in *Pyricularia* (unpublished data), these presumably were cases of cross-infection. Isolates from *Cenchrus, Digitaria*, and *Pennisetum* that failed to yield CH7BAC9 amplicons were characterized by amplifying and sequencing the MPG1 locus. Phylogenetic analysis of the resulting data, along with sequences from a number of reference isolates including *P. oryzae*, revealed that all of the isolates from *Digitaria* (n = 45) were *P. grisea*, while those from *Cenchrus* (n = 23), and the one from *Pennisetum* were *P. pennisetigena* (Fig. 2B). The MPG1 allele in the isolate from *Pennisetum* had the *P. pennisetigena* genotype. *P. pennisetigena* is named because the type isolate came from *Pennisetum* (Klaubauf et al. 2014). However, the present data indicate that *Cenchrus* is also a canonical host for this species.

### GBS confirmed that most *P. oryzae* lineages are host-specialized

The combined results of the MoT3/C17 assays and CH7BAC9 sequencing identified very few PoT isolates on endemic grasses and, conversely, very few grass-adapted isolates on wheat (Table 6). However, because a small proportion of PoT isolates are known to lack MoT3, we considered it important to rule out the possibility that shifts in the PoT population had produced isolates that lack C17, or both MoT3 and C17. Therefore, to validate the lineage assignments made with MoT3, C17, and CH7BAC9, we used “MonsterPlex” - Floodlight Genomics LLC’s variation of the Hi-Plex2 assay (Hammet et al. 2019) - to perform genotyping-by-sequencing (GBS) on a selection of isolates from wheat (n = 66), *Urochloa* (n = 38), *Eleusine* (n = 6), *Hordeum* (n = 3), *Melinis* (n = 6) and *Panicum* (n = 11). We then examined their phylogenetic relationships to *in silico*-mined genotypes from a set of 232 reference isolates whose lineage affiliations were already well established (e.g., Gladieux et al. 2018).

**TABLE 6.** Frequencies of positive (indicating a *Pyricularia oryzae* Triticum lineage - PoT) and negative (indicating a non-PoT lineage) amplifications of the C17 primer (Thierry et al. 2020) for a set of 487 *Pyricularia* isolates obtained from leaves of grass plants located away or nearby wheat fields or from leaves or heads of wheat plants displaying the typical blast symptoms across several locations in MG state, Brazil.

| Host | Proximity to wheat | C17+ (PoT)<br>Count (%) | C17- (non-PoT)<br>Count (%) | Sum |
| --- | --- | --- | --- | --- |
| Wheat | - | 322 (97.9%) | 7 (2.1%) | 329 |
| Grass | Nerby (< 1 km) | 7 (10.6%) | 59 (89.3%) | 66 |
|  | Away (> 30km) | 2 (2.1%) | 90 (98.1%) | 92 |
| Sum |  | 331 | 156 | 487 |

Although the multiplex assay was originally designed to target only 84 SNPs, we identified a total of 228 variant sites within the targeted loci. Together, these SNPs were capable of resolving all 34 of the PoT haplotypes known to exist prior to this study (Rahnama et al. 2021). Sixty-two of the 64 MG isolates identified as PoT using PCR-based diagnostics grouped with one of the two established PoT/PoL clades (Fig. 3). For the remaining pair of isolates, the GBS data revealed that they had been mis-characterized as PoT based on their MoT3/C17 amplification profiles (MoT3^-^/C17^+^). One was phylogenetically related to *Urochloa* pathogens (PoU3), and the other grouped with isolates from *Melinis* (PoM) (Fig. 3). These isolates represent the first false positives to have come from C17 diagnostics. Also analyzed were two isolates from wheat for which the original PCR tests failed altogether. One was found to be a PoU3 member, and the other grouped with other *Urochloa* pathogens in the PoU4 clade, which is phylogenetically related to *Panicum* pathogens (PoP).

**Fig. 3.**
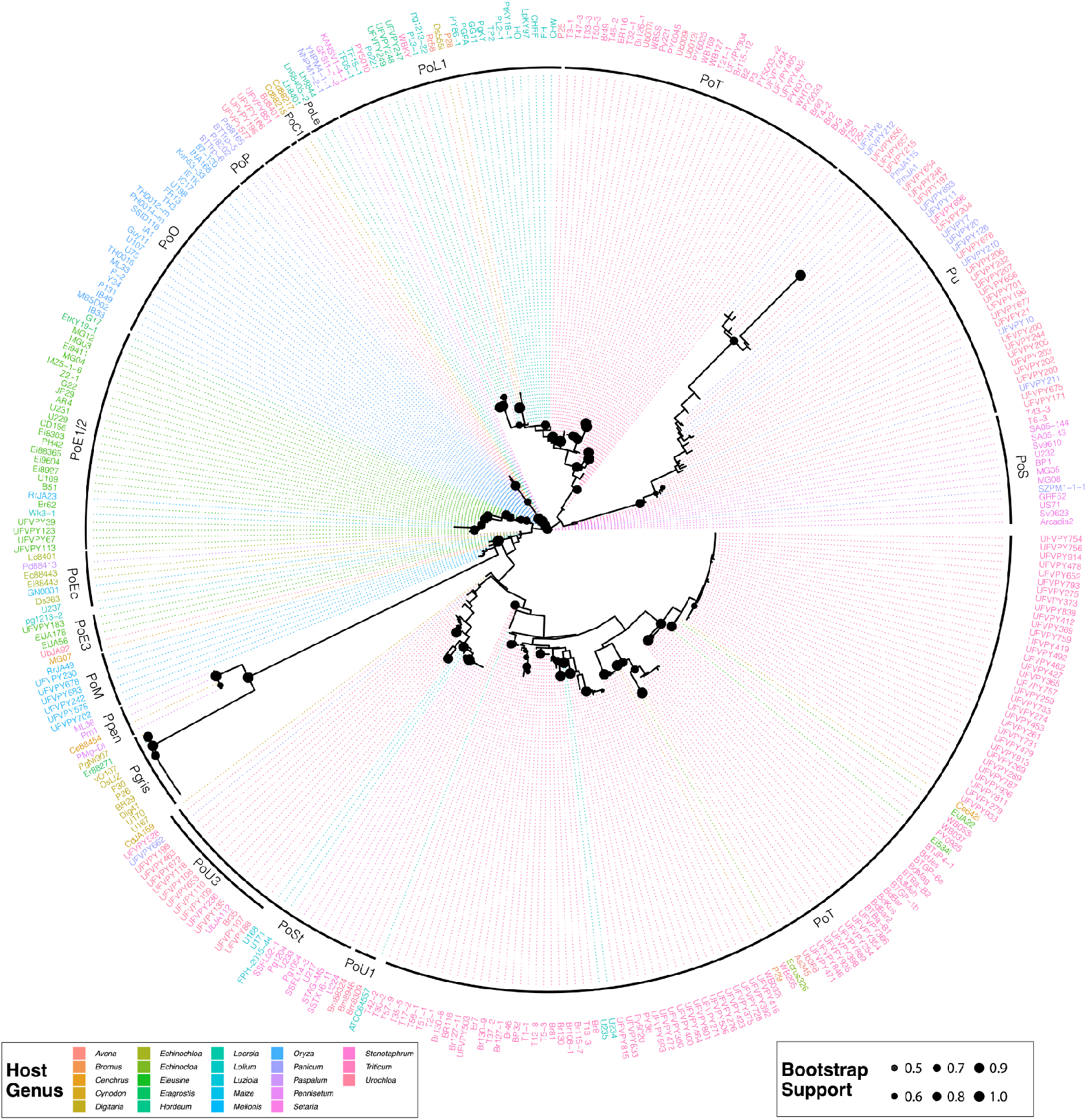
Maximum Likelihood tree showing phylogenetic placement of *Pyricularia* isolates from MG as determined using “MonsterPlex” genotyping-by-sequencing. Isolate names are colored according to host-of-origin and those from MG are identified with a UVFPY prefix. Phylogenetic lineages are highlighted with black lines and are labeled according to the primary host (as noted in Figure 2). PoU3 forms a subgroup of PoSt but is labeled separately to emphasize that it constitutes a key, *Urochloa*-infecting population. Nodes with bootstrap support ≥ 0.5 are highlighted with proportionally-sized circles.

GBS was also performed for MoT3^-^ and/or C17^-^ isolates from non-wheat hosts (*Eleusine, Melinis, Panicum*, and *Urochloa*) to test for possible cross-infections by PoT members with atypical genotypes. No such evidence was obtained because all isolates analyzed grouped outside of the PoT clades. The only potential cross-infections identified involved isolates from *Hordeum vulgare* (UFVPY247, 248, 249), which grouped with PoL1. Isolates from *Eleusine* and *Melinis* grouped strictly according to their respective hosts of origin, with the *Eleusine* pathogens belong to PoE1/2 (n = 4) and PoE3 (n = 1), and the *Melinis* isolates to PoM (n = 6). Only eleven of the 38 isolates from *Urochloa* belonged to a previously defined PoU lineage - this being PoU3. The remainder grouped in two novel *Urochloa*-associated clades, one related to torpedograss (*Panicum repens*) pathogens (PoU4, n = 3 isolates), and the other, a lineage that appears to fall under the umbrella of the sister species, *P. urashimae* (Pu, n = 24) because it houses PmJA1 and PmJA115 (Fig. 3), and these isolates possess Pu alleles for a number of reference genes. It should be noted that there were a large number of missing datapoints for isolates within the Pu lineage, presumably due to significant sequence divergence at the target loci affecting primer binding (∼10%). However, there was also significant sequence divergence among the successfully amplified sites.

### Identification of false positives for the MoT3 and C17 diagnostic markers

A large fraction of the isolates from *Urochloa* and *P. maximum* exhibited a MoT3^+^/C17^-^ genotype, which implied that these isolates are false positives for MoT3. This was confirmed using GBS (and genome sequencing, see Farman et al. 2022) which revealed that all of the MoT3^+^ *Urochloa* pathogens are PoU3 and the positive *Panicum* pathogens belong to the *P. urashimae* lineage (Fig. 3). Conversely, isolate UFVPY183 from *Eleusine* (PoE3) and UFVPY578 from *Melinis* (PoM) reproducibly tested positive for C17 and negative for MoT3 and, therefore, are the first examples of non-PoT isolates that have given positive results for C17.

### *Triticum-Urochloa* specificity observed in the field may be due to inherent differences in infection capability on canonical versus non-canonical hosts

Molecular analysis of field isolates collected from wheat and *Urochloa* revealed that cross-infection between the two hosts is uncommon. To explore whether this is due to inherent differences in relative aggressiveness toward the respective hosts, we performed reciprocal infection assays. In a first experiment, 14 PoT isolates (11 from wheat and three from *Urochloa*) and six isolates that were obtained from *Urochloa* (three from the PoU3 lineage; two from Pu and one from PoU4 - hereafter non-PoT) were inoculated on leaves of two wheat cultivars (Guamirim and BRS18-Terena) and leaves of one signalgrass cultivar (Marandu). In general, the isolates within each lineage were consistently and significantly more aggressive on their primary host of origin (with the exception of the PoTs obtained from *Urochloa)* than on the alternative host, although there were differences among individual isolates (Figs. 4, 5, 6 and 7).

**Fig. 4.**
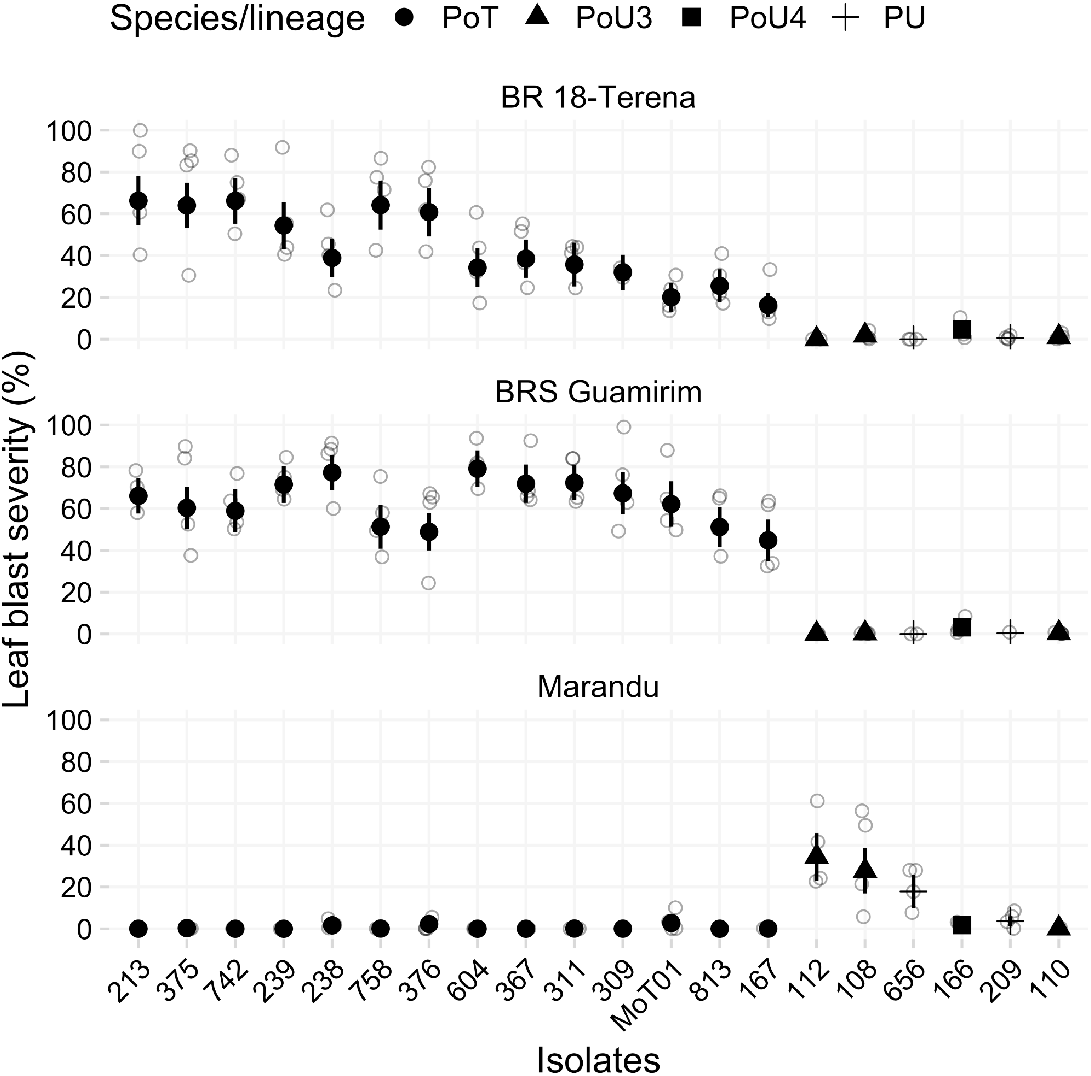
Aggressiveness, expressed as percentage leaf area affected (% severity), evaluated in replicated greenhouse experiments (foliar inoculations on two wheat cv. [BR18 Terena and Guamirim and one signalgrass cv [Marandu]), for a set of 19 isolates (10 from wheat [*Triticum aestivum*], and 9 [six non-PoT] and three PoT = 742, 758 and 213] from signalgrass [*Urochloa* spp.]) of *Pyricularia oryzae* collected in MG state, Brazil and which showed a positive (indicating Triticum lineage) or a negative (indicating non-Triticum lineage) reaction when screened moleculary using the C17 primer set (Thierry et al. 2020). The non-PoT isolates were further identified with genotype by sequencing: isolates 108, 110 and 112 = *Pyricularia oryzae* lineage *Urochloa3*; 166 = *P. oryzae* lineage *Urochloa4;* 209 and 656 = *P. urashimae*. The MoT01 strain is PoT used as a reference for an aggressive isolate collected in Passo Fundo, Brazil, and used (coded as 16MoT001) in screening for host resistance studies (Cruppe et al. 2020). The empty circles represent values of replicates (two experiments combined), the symbols represent the mean values and the error bar is the 95% confidence limit.

**Fig. 5.**
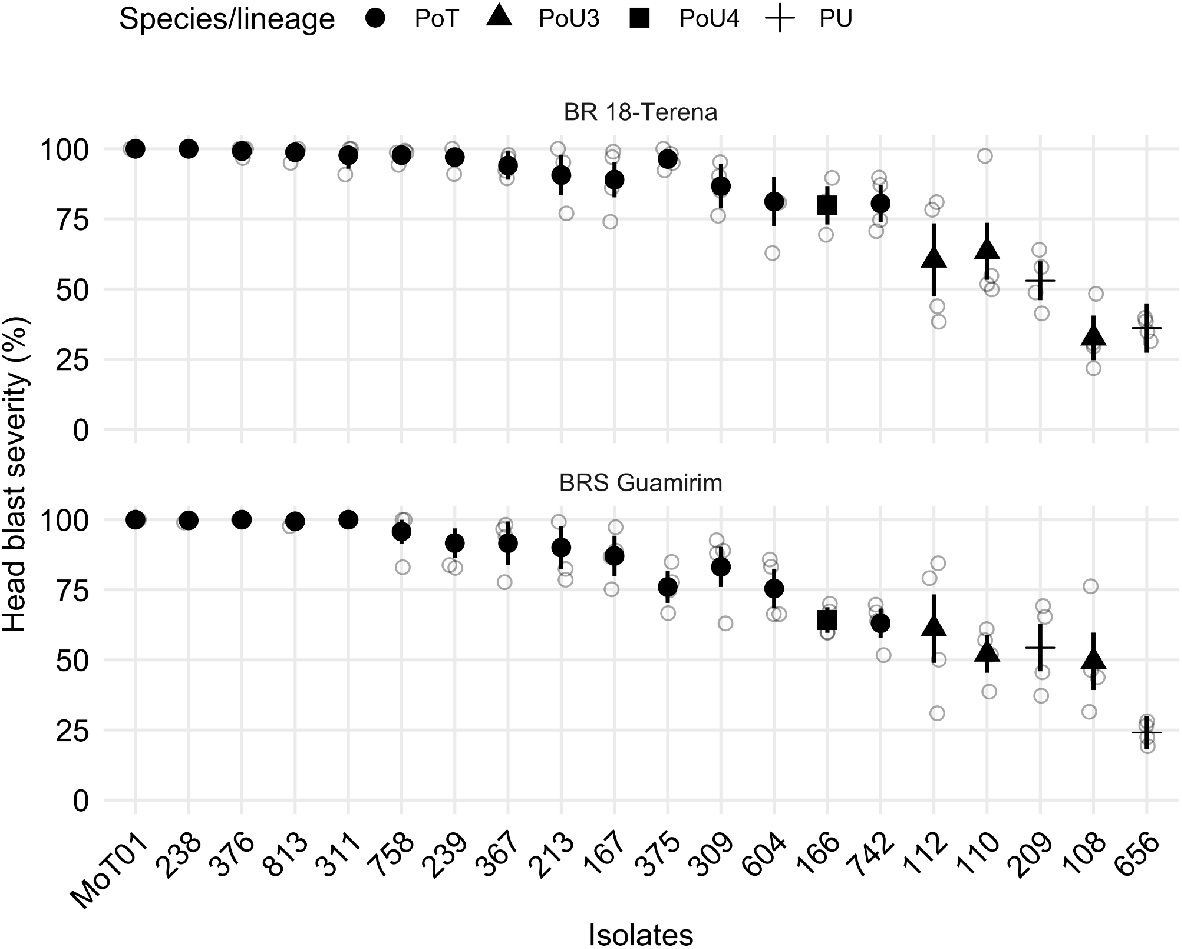
Aggressiveness, expressed as percentage of affected spikelets (% severity), evaluated in replicated greenhouse experiments (head inoculations on two wheat cv., BR18 Terena and BRS Guamirim) for a set of 19 isolates (10 from wheat [*Triticum aestivum*], and 9 [six non-PoT and three PoT = 742, 758 and 213] from signalgrass [*Urochloa* spp.]) of *Pyricularia oryzae* collected in MG state, Brazil and which showed a positive (indicating Triticum lineage) or a negative (indicating non-Triticum lineage) reaction when screened moleculary using the C17 primer set (Thierry et al. 2020). The non-POT isolates were further identified with genotype by sequencing: isolates 108, 110 and 112 = *Pyricularia oryzae* lineage *Urochloa3*; 166 = *P. oryzae* lineage *Urochloa4;* 209 and 656 = *P. urashimae*. The MoT01 strain is a reference for an aggressive isolate collected in Passo Fundo, Brazil, and used (coded as 16MoT001) in screening for host resistance studies (Cruppe et al. 2020). The empty circles represent values of replicates (two experiments combined), the symbols represent the mean values and the error bar is the 95% confidence limit.

**Fig. 6.**
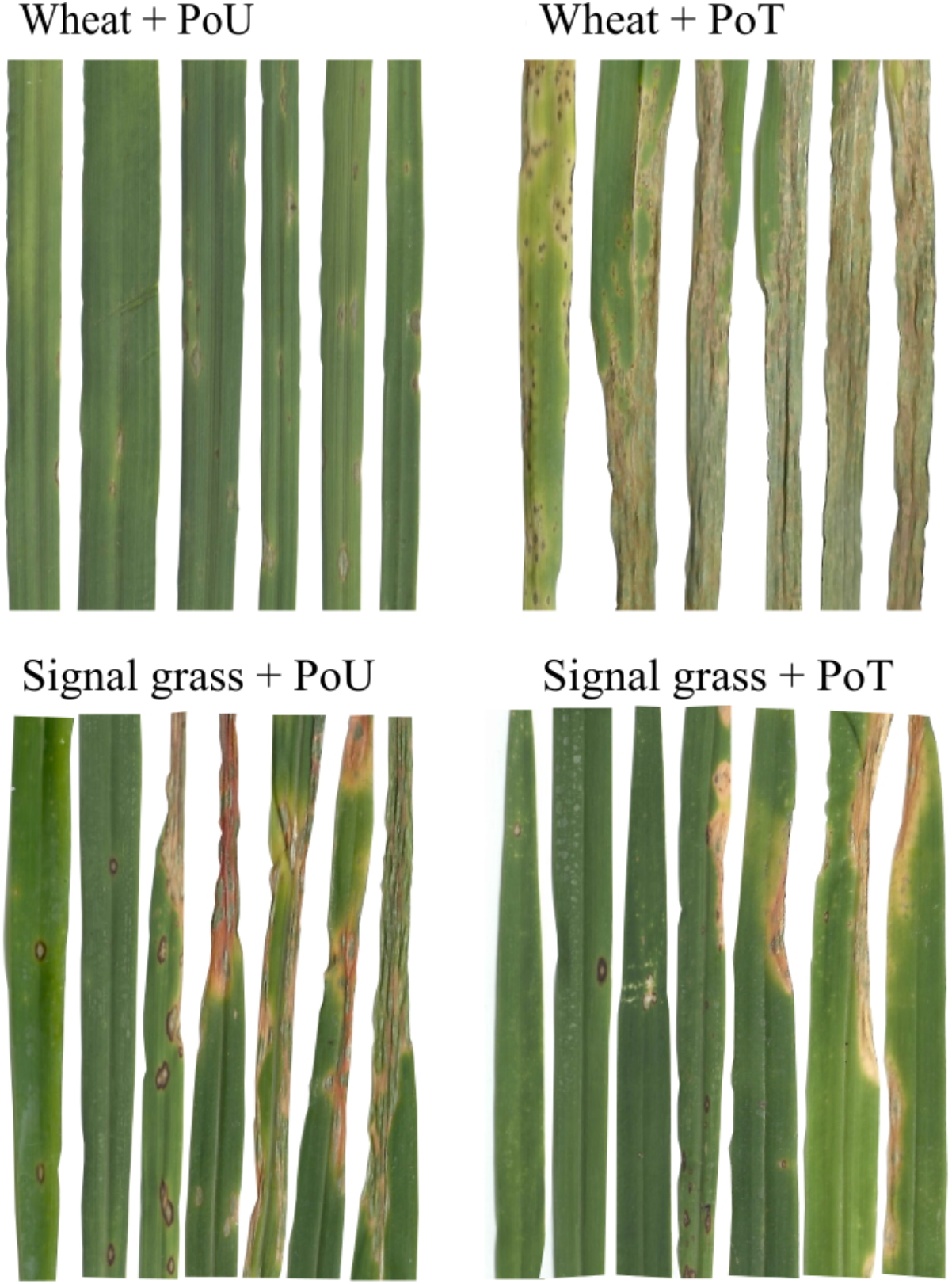
Leaf blast disease symptoms resulting from cross-inoculations of *Pyricularia oryzae* lineages (PoT = *Triticum* lineage isolated from wheat; PoU3 = *Urochloa* lineage 3 isolated from signalgrass) on *Triticum aestivum* (cvs. BRS Guamirim) and *Urochloa brizantha* (cv. Marandu) under greenhouse conditions.

**Fig. 7.**
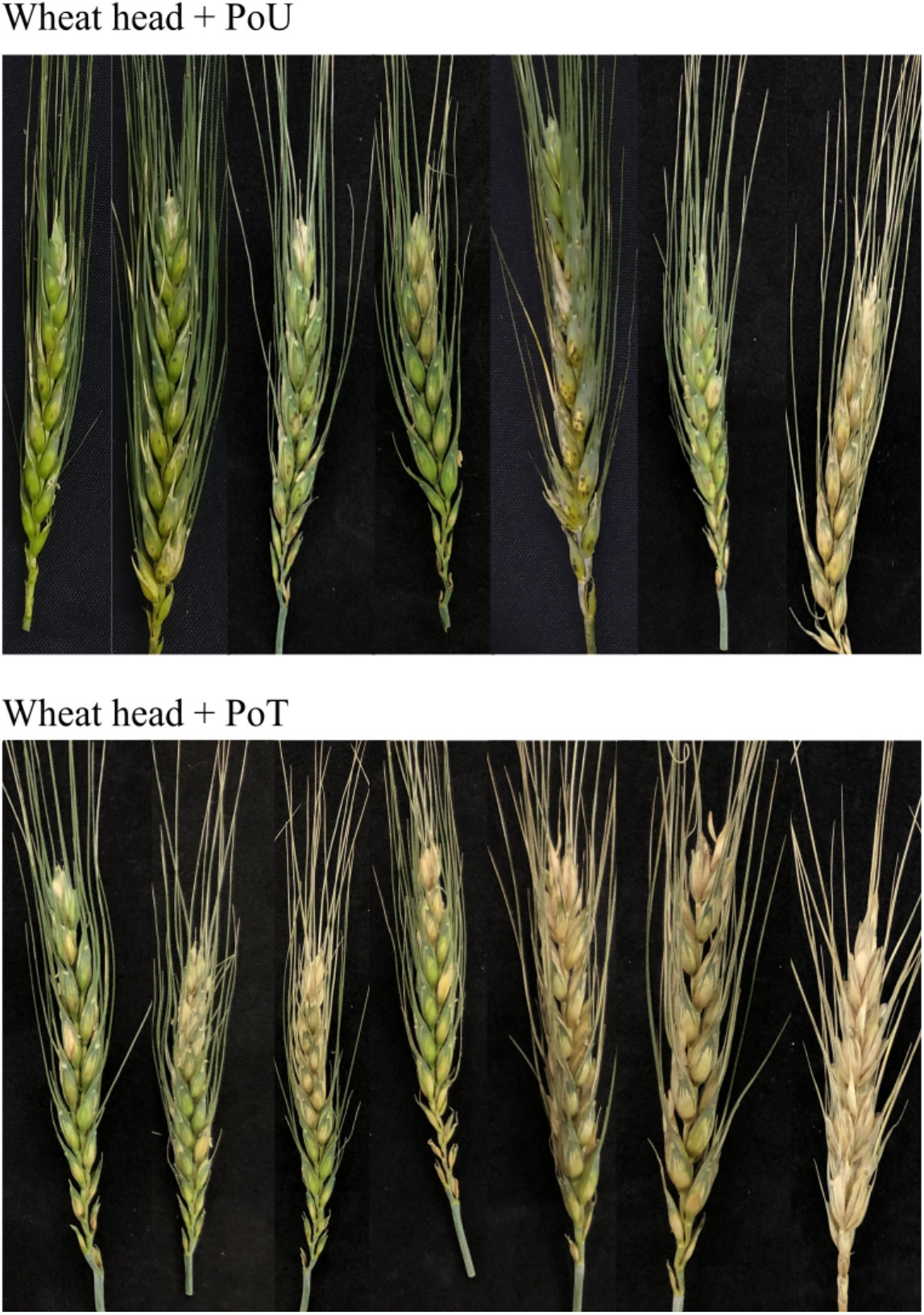
Wheat head blast symptoms resulting from inoculation of *Pyricularia oryzae* lineages (PoT and PoU3) on *Triticum aestivum* (BRS Guamirim) under greenhouse conditions. PoT = *Triticum* lineage, isolated from wheat; PoU3 = *Urochloa* lineage 3 isolated from signalgrass.

On average, severity on the leaves induced by the PoT isolates ranged from 20 to 70% (mean = 44.1%) on BR18 Terena wheat and from 40 to 80% (mean = 63.1%) on BRS Guamirim wheat, across isolates. Severity induced by the non-PoT isolates on leaves of wheat was only 1.39% and 0.73% on BR18 Terena and BRS Guamirim, respectively. Mean severity induced by PoT and non-PoT on Marandu signalgrass was 0.51% and 14.26%, respectively (Fig. 4). In the separate experiment, wheat heads of the same two cultivars were inoculated with the same set of PoT and non-PoT isolates.

The PoT isolates were generally more aggressive than the non-PoT isolates (Fig. 5 and 7). The percent of infected spikelets were, on average, 89% and 93% on BRS Guamirim and BR18 Terena, respectively, when challenged with PoT isolates, including those three isolates obtained from *Urochloa* (Fig. 5). Contrarily, percent infected spikelets by the non-PoT isolates were on average 54.2% and 50.9% across the isolates (Fig. 5). It is worth noting that lesions caused by the non-PoT isolates on the affected spikelet were small and scattered, not affecting the entire spikelet (Fig. 7). On the other hand, most PoT isolates were highly aggressive, producing the typical bleaching of the affected spikelets (Fig. 7).

## Discussion

Over four separate sampling trips spanning two years (prior to and during the wheat growing season), we generated a comprehensive collection of *Pyricularia* isolates obtained from blast lesions on endemic grasses grown near to or away from wheat fields. The grass genera from which we recovered non-PoT *Pyricularia* were *Cenchrus, Cynodon, Digitaria, Eleusine, Hordeum, Melinis, Panicum, Pennisetum*, and *Urochloa*. At the same time, we established the first extensive collection of several hundreds of isolates obtained from wheat cultivated in both southern and western regions of MG, Brazil.

A large majority of isolates could be reliably identified down to species/lineage through amplification/sequencing of just three PCR-based markers. Successful amplification of CH7BAC9 by itself distinguished *P. oryzae* from the other species, and positive amplification for both MoT3 and C17 (Pieck et al. 2017; Thierry et al. 2020) proved to be definitive for PoT. Amplification of either MoT3 or C17 alone, however, yielded equivocal results. Although MoT3 showed early promise as a PoT diagnostic, occasional exceptions (false positives/negatives) have been reported (Pieck et al. 2017; Yasuhara-Bell et al. 2018). In the past, the most common exceptions involved wheat-infecting members of the related PoL1 lineage which are MoT3^-^/C17^-^. We found that 2.5% (9/394) of the wheat blast isolates from MG wheat fell into this category. More concerningly, however, we found an extremely high frequency of false positives, with 41% (64/157) of non-PoT isolates yielding a MoT3^+^ reaction. This result is attributable to the fact that PoT/PoL1 have hybrid genomes, and the MoT3 locus was acquired from the PoU3 lineage (Rahnama et al. 2021), which is abundantly represented among *P. oryzae* isolates from *Urochloa*. Additionally, a highly similar, and amplifiable, MoT3 sequence was ubiquitously present in *P. urashimae* isolates from *Panicum maximum* (14/14) (Table S1). Consequently, assays that survey MoT3 alone are unreliable for PoT detection, which throws into question conclusions from a recent study which used MoT3 amplification to assess the presence of PoT on Brazilian grasses, including *Urochloa* spp. (Maciel et al. 2023). Because all MoT3^+^ isolates in that study came from *Urochloa* (13/58; 22.4%), it is quite possible that none of the isolates sampled in that study were PoT, especially considering the low frequency of PoT we found on *Urochloa*. It would, therefore, be instructive to reassess PoT prevalence after performing C17 assays.

Here, it should also be stressed that the specificity of the C17 assay was also not perfect. Although C17 yielded positive results for the nine PoT isolates that were MoT3^-^, we recorded the first examples of false positives in isolates from *Eleusine* (PoE3) and *Melinis* (PoM). This, again, is not surprising because PoT inherited the C17 locus from another - in this case, unknown - *P. oryzae* lineage (PoX) (Rahnama et al 2022), which means that this locus, too, will yield false positive results when isolates from the relevant host(s) are sampled. Therefore, for reliable PoT detection, at a minimum we recommend that both MoT3 and C17 be surveyed in parallel. Finally, it is also important to note that the current formulation of the MonsterPlex assay, while being highly effective at lineage assignment for most isolates, and identifying new lineages, is also not capable of positively identifying PoT. This is because several known PoT isolates group with PoL1, while others such as PoT6 and PoT29, group with neither PoT, nor PoL1 (Fig. 3). This is not surprising because the assay was originally designed with the main goal of distinguishing the B71 lineage from all other PoT (Tembo et al. 2021).

Overall, among the 572 *Pyricularia* spp. isolates, *P. oryzae* dominated the collection (87%) and this likely reflected the fact that other *Pyricularia* were much more host-restricted, with *P. pennisetigena* being found almost exclusively on *Cenchrus, P. grisea* on *Digitaria*, and *P. urashimae* on *Panicum* and *Urochloa*. By way of contrast, *P. oryzae* was recovered from all the genera that yielded *Pyricularia*, except *Cenchrus*; and accounted for all but two of the isolates sampled from wheat heads (n = 333). Thus, species outside of the *P. oryzae* clade are very unlikely to cause wheat blast. This is contrary to what has been suggested following controlled environment studies, which reported pathogenicity and high aggressiveness of *P. pennisetigena* and *P. zingibericola* - from grasses in Brazil - to Anahuac 75, a wheat cultivar regarded as universally susceptible cultivar to PoT (Reges et al. 2016).

The *P. oryzae* isolates we collected from non-wheat/*Lolium* hosts grouped into nine distinct lineages/species variously specialized on eight different grass genera, most of which were previously known hosts, including *Cynodon, Echinochloa, Eleusine, Urochloa* (Borromeo et al. 1993), and *Hordeum* (Urashima et al. 2004). Although we identified members of previously known lineages (PoE1/2) in association with the expected hosts, a majority of isolates belonged to new phylogenetic lineages. These also showed evidence of host-specialization because constituent members were usually isolated from the same genus/species. These included PoE3 specialized on *Eleusine*, PoM (from *Melinis*), PoU3 and PoU4 (both on *Urochloa*). One additional lineage was identified (PoC2 from *Cynodon*) but was only represented by one isolate, so its host-specialization status is unclear.

We also report *P. maximum* as a new host for *P. urashimae* (Pu), the type isolate of which originally came from *Urochloa brizantha* (Crous et al. 2016). This seems to be quite a specific interaction from *P. maximum*’s perspective because 12/16 isolates from this grass were in the Pu phylogenetic group, with limited cross-infection by PoM, and PoT having been observed. However, most Pu members came from *Urochloa*, indicating that the lineage has dual specificity. This property might be partially explained by the GBS data, which suggests that the Pu species is highly diverse, such that it too, like *P. oryzae*, might comprise a number of genetically-distinct sub-lineages, with some being specialized on *P. maximum*, and others on *Urochloa*.

Although our primary motivation for sampling wheat blast was to examine the extent of cross-infection by grass-adapted lineages, the GBS data also provided key insights into the MG wheat blast population. A recent phylogenomic analysis revealed that wheat blast and gray leaf spot co-evolved very recently through a series of admixtures involving *P. oryzae* isolates from five different host-specialized populations. This resulted in a set of distinct chromosomal haplotypes, that are defined by the specific chromosome segments they inherited from the various donor isolates (Rahnama et al. 2021). To date, 37 distinct haplotypes have been found on wheat (34 PoT; 3 PoL1). The MG wheat blast isolates defined eight phylogenetic groups, none of which perfectly matched the known haplotypes; and because so few mutations have arisen since wheat blast/gray leaf spot evolved (Rahnama et al 2022), the MG blast population most likely have new chromosomal haplotypes. Given that all of the PoT isolates analyzed previously came from other states, namely Paraná, Rio Grande do Sul, Mato Grosso do Sul, Goiás, and São Paulo, this suggests that there may be regional differences in the genetic composition of the South American wheat blast population.

We found that the vast majority of *Pyricularia* isolates collected from endemic grasses in MG state were genetically distinct from PoT, with most belonging to phylogenetic groups consistent with their host-of-origin. This is in striking contrast with the findings of prior studies of *Pyricularia* collected from grasses in Brazil, where all but one of the characterized isolates were members of the PoT or PoL1 populations. We conclude that these earlier studies made the mistake of only sampling infected weeds beneath, or immediately adjacent to, infected wheat plants, as implied by the GPS coordinates provided in the relevant reports (Maciel et al. 2014; Castroagudín et al. 2016). Consistent with this interpretation, most of the weeds we found to be harboring PoT were collected near to heavily infected wheat plots. There appears to be no obvious pattern to PoT’s cross-infectivity, because we identified it on six different host genera. Together with the data from prior studies (Castroagudín et al. 2016; Maciel et al. 2014), this expands the list of alternative hosts for PoT to ten (*Bromus, Cenchrus, Digitaria, Echinochloa, Eleusine, Lolium, Melinis, Panicum, Pennisetum*, and *Urochloa*). Further, if we include PoL1 isolates based on the fact that some of its members can also infect wheat, this also adds *Avena*, and *Hordeum* as potential surrogate wheat blast hosts.

The discovery of seven non-PoT/PoL isolates on wheat was rather surprising because, with the exception of PoL1 lineage members, cross-infection of wheat by other host-adapted forms of *P. oryzae* has never been shown beforehand. Four of the isolates were collected in the same wheat field in the Triangulo Mineiro region, along with 40 PoT strains. It is possible that we were successful in identifying these novel cases due to the repeated sampling from the same field, and the fact that we specifically screened for fungal isolates that were MoT3/C17^-^.

Here, we greatly expand on understanding of the *Pyricularia* populations colonizing endemic grasses in Brazil, especially when we consider that prior efforts to characterize grass-infecting populations in Brazil (Castroagudín et al. 2017; Castroagudín et al. 2016; Ceresini et al. 2018, 2019) ended up sampling just one isolate from a grass-adapted lineage (Farman et al, 202). That isolate, Ds555i (a.k.a. 12.1.555i, from *Digitaria*), is a member of the PoEc (*Echinocloa*) lineage which distinguishes itself by the absence of host-specialization among its constituent members. Interestingly, we found two PoEc members on a previously unknown host of this lineage - *Eleusine*. As a rule, we found cross-infections to be fairly uncommon in MG, with most *Pyricularia* populations exhibiting significant host-specialization and, although members of non-adapted lineages were routinely recovered from most of the sampled genera, the isolates found on non-canonical hosts were fairly evenly distributed among the different lineages. This finding, along with the discovery that certain host genera are susceptible to multiple genetically-distinct lineages (e.g. *Cynodon, Eleusine, Hordeum, Panicum, Urochloa*) implies that host-specificity barriers to *P. oryzae* are somewhat fluid and, therefore, reinforces the notion that most lineages are “host-adapted” or “host-specialized,” as opposed to “host-specific.”

A main focus of our study was to characterize the fungal population(s) found on *Urochloa* because we suspected that prior studies implicating this host as a central player in the evolution, inoculum development and epidemic spread of wheat blast (Ceresini et al. 2018, 2019; Maciel et al. 2014; Stukenbrock and McDonald, 2008) were incorrect due to flaws in both sampling and phylogenetic inference (see Farman et al. 2022). Previous studies only sampled two *P. oryzae* lineages from *Urochloa* - PoT and PoL1 (Castroagudín et al. 2017; Castroagudín et al. 2016; Ceresini et al. 2018, 2019). Here, we identified an additional seven lineages on the genus (PoU1, PoU2, PoU3, PoU4, PoE3, PoL, PoM, PoSt, and PoT), as well as *P. urashimae*. At first, this might imply that *Urochloa* is a “universally susceptible” host. However, most isolates were placed in the *Urochloa*-adapted lineages, PoU3 and PoU4, or the highly diverse *P. urashimae* (Pu) clade, which is dually specialized on *Urochloa* and *P. maximum*. Thus, the small number of isolates from various other lineages found on *Urochloa*, probably reflects a low level of inherent, base-line cross-infectivity across the species.

We found that PoT was rarely recovered from infected *Urochloa* plants sampled away from wheat plots with only one out of 67 isolates being a PoT member. The low frequency of wheat <-> *Urochloa* cross-infection found in nature seems to be correlated with innate compatibility differences because, using cross-inoculation experiments, we found that PoT members were consistently more aggressive on wheat leaves, and the isolates from *Urochloa* (PoU3/4 and Pu) were more aggressive on signalgrass. Thus, our findings hold to the general pattern that PoT is usually more aggressive on wheat, while non-PoT/non-*P. oryzae* isolates tend to be less aggressive, even under favorable, controlled environments (Chung et al. 2020; Kato et al. 2000; Reges et al. 2016, 2019). And, even though the *Urochloa* pathogens showed incidences on the spikelets up to 50%, after spraying spikelets, the percent diseased area was apparently rather low, and the symptoms were mild, consisting of small, reddish-brown to dark-gray spots, or even hypersensitive reactions. This was in striking contrast to the symptoms caused by PoT isolates, which were characterized by bleaching of the heads. It seems doubtful that inconspicuous lesions on spikes caused by non-PoT isolates are likely to cause significant yield loss, although this should be further confirmed using polycyclic infection assays. It should also be noted that *Urochloa* leaves collected from the field often showed an abundance of blast-like symptoms but the vast majority of lesions failed to produce the profuse sporulation characteristic of *Pyricularia* infection after overnight humidification. True blast infections typically show rapid and abundant sporulation from all lesions and, therefore, it appears that not only was PoT rarely recovered from *Urochloa* but the incidence of blast was also a lot lower than was initially apparent based on macroscopic symptoms.

Prior studies have implied that cross infection of *Urochloa* by PoT is significant and widespread (Castroagudín et al. 2017; Ceresini et al. 2018, 2019; Maciel et al. 2023), and a major concern for wheat blast management. Overall, our data challenge this idea because most *Pyricularia* isolates from signalgrasses - even ones collected in proximity to wheat fields - belonged to *Urochloa*-specific lineages; and members of these lineages were recovered from wheat even less frequently than was PoT from *Urochloa*. In addition, blast-like lesions were rarely observed on signalgrasses growing at remote distances from wheat fields, and were sometimes even hard to find on signalgrass plants immediately adjacent to devastated wheat (personal observations). Thus, we feel we can propose an equally viable hypothesis that infected wheat more often serves as a source of inoculum for the occasional cross-infection of nearby *Urochloa*. Of course, we cannot rule out the possibility that the low prevalence of PoT on *Urochloa*, and the low sporulation capacity, might still be sufficient to produce an inoculum reservoir to trigger seasonal epidemics, and facilitate long-range movement. However, it should be noted that PoT was found as often on other weedy grasses, as it was on *Urochloa* and, therefore, the proportional contributions of these different grasses, if any, to wheat blast epidemiology remains an open question.

## Acknowledgements

This work was supported by the United States Department of Agriculture, Agriculture and Food Research Initiative grant 2013-68004-20378, multistate project NE1602; Agricultural Research Service project 8044-22000-046-00D; Hatch project KY012037; the National Science Foundation, MCB-1716491; and the University of Kentucky College of Agriculture Food and the Environment. Emerson M. Del Ponte was supported by the National Council for Scientific and Technological Development (CNPq) through a Productivity Research Fellowship (PQ) project 310208/2019-0, and through research grants provided by FAPEMIG. João P. Ascari and Luis I. Cazón were supported by CNPq through doctoral scholarships.

## Supplementary materials

**Fig. S1.**
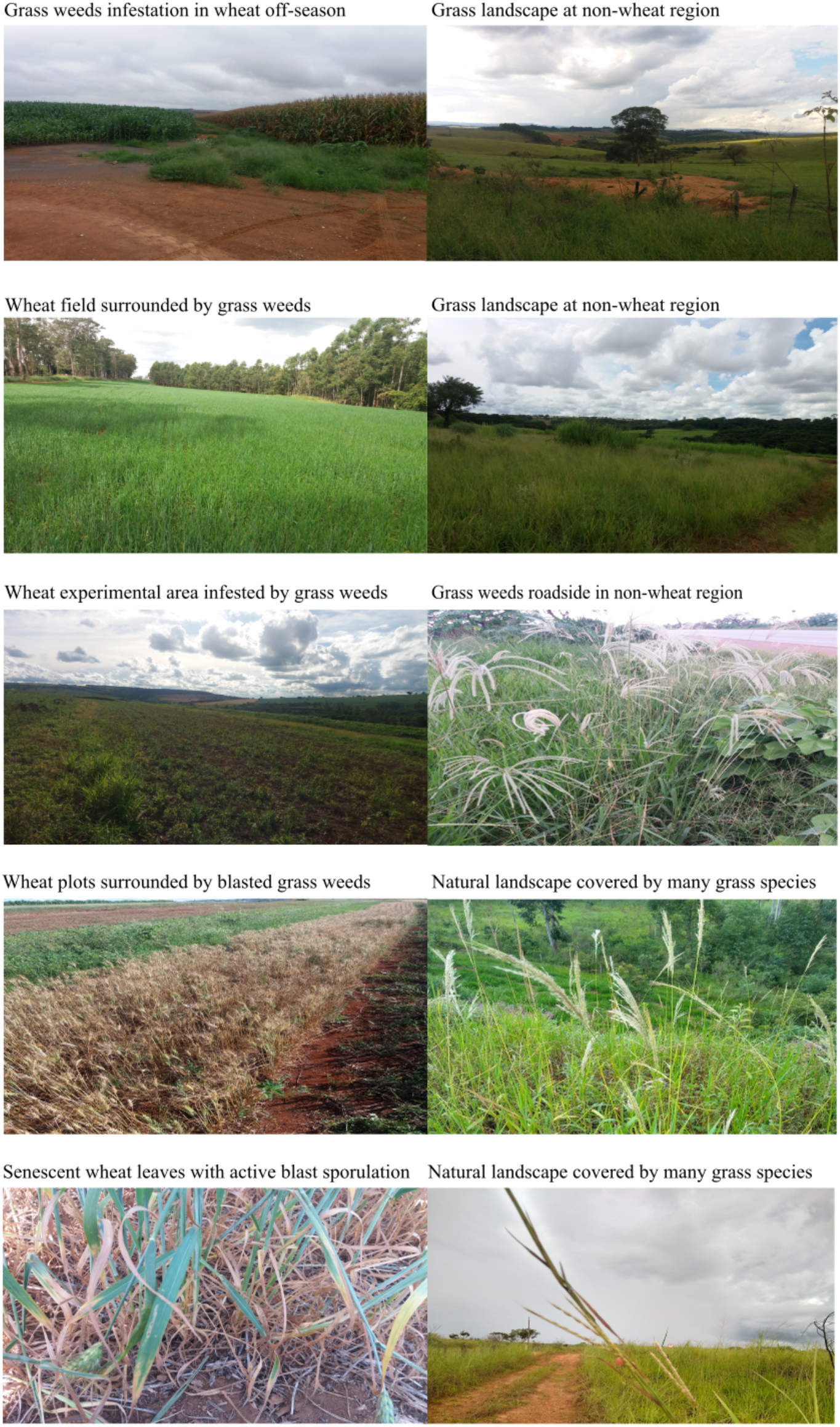
Images of the natural landscape and wheat commercial fields where grass weeds and wheat were collected during surveys conducted during the pre-season summer (February) and within-season fall (May) in 2018 and 2019, at Triângulo Mineiro and Centro-Sul de Minas, Brazil.

**Fig. S2.**
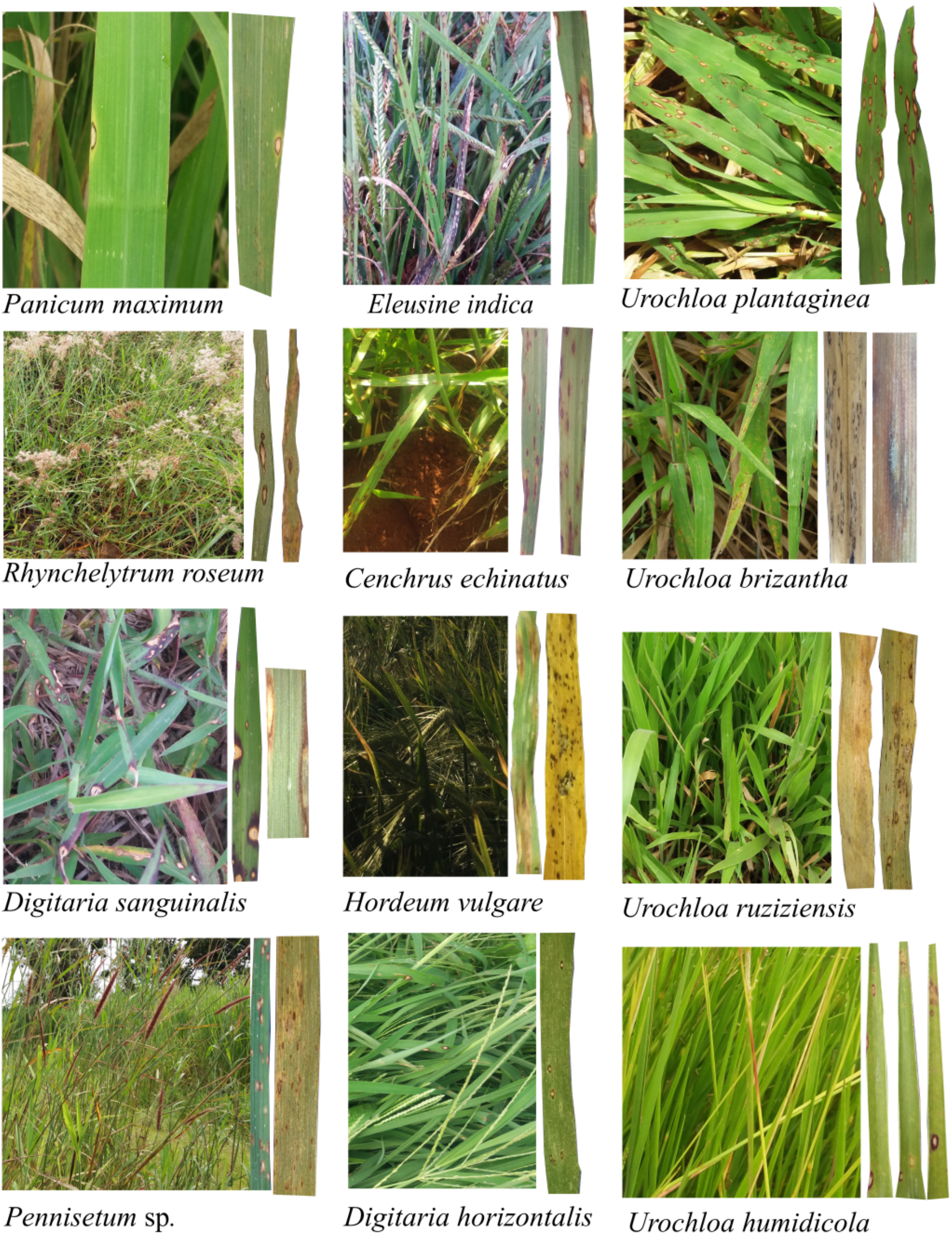
Details of symptoms on full plants and leaves of species within Poaceae exhibiting typical blast symptoms. The surveys were conducted twice in 2018 and also 2019, at Triângulo Mineiro and Centro-Sul de MG, Brazil. Plant identification was made at the species level, whenever possible, based on adult specimen morphological characteristics (Crispim and Branco 2002; Lorenzi 2014).

